# Correcting for Global Synonymous Selection Improves the Accuracy of Episodic Positive Selection Inference

**DOI:** 10.64898/2026.06.02.729680

**Authors:** Hannah Verdonk, Alyssa Pivirotto, Jody Hey, Sergei L Kosakovsky Pond

## Abstract

The ratio of nonsynonymous to synonymous substitution rates (*ω*) constitutes a fundamental parameter for inferring adaptive protein evolution, predicated upon the assumption that synonymous substitutions are selectively inert. This premise, however, is increasingly untenable given evidence of selection acting on synonymous substitutions, driven by various biological processes such as translational efficiency and mRNA stability. In this study, we demonstrate that unmodelled synonymous selection introduces substantial bias into *ω* estimation, resulting in elevated false positive rates in tests for positive selection. To rectify this, we present BUSTED+S+MSS, a statistical framework incorporating Multiclass Synonymous Substitution (MSS) models into BUSTED, a method for detecting episodic selection. By partitioning synonymous codons into empirically derived rate classes, this approach accounts for global synonymous constraints. Application to five diverse clades—*Drosophila, Caenorhabditis, Enterobacteria, Saccharomyces*, and Primates—reveals that the inclusion of MSS components consistently improves model fit and reduces the proportion of genes inferred to be under positive selection. In *Enterobacteria*, genes retaining significance under the corrected model exhibit weaker constraint on synonymous substitutions (*dSs*), consistent with the hypothesis that unmodelled purifying selection drives spurious signals of adaptation. Furthermore, an information-theoretic analysis indicates that whilst site-specific variation (SRV) provides the primary correction, global synonymous rate variation (MSS) contributes a distinct second-order correction. In highly divergent alignments, these signals act in concert to improve model fit. The BUSTED+S+MSS framework, especially when coupled with an “error-sink” to absorb alignment artifacts, thus offers a computationally feasible means to disentangle adaptive nonsynonymous substitution from the confounding effects of synonymous constraint.

## 1 Introduction

The inference of natural selection acting upon protein-coding genes constitutes a primary objective in evolutionary biology, elucidating the functional constraints and adaptive innovations that shape genomic diversity. Central to this endeavour are codon-based models of sequence evolution, which estimate the ratio of non-synonymous to synonymous substitution rates (*ω*) ((Muse and Gaut, 1994; Goldman and Yang, 1994; Yang et al., 2000; Pond et al., 2005). An *ω* value exceeding one is interpreted as evidence of positive diversifying selection, whilst values below one indicate purifying selection.

The validity of this metric is predicated upon the assumption that synonymous substitutions are selectively inert and thus provide an unbiased baseline for the rate of mutation. However, this assumption is increasingly challenged by evidence that synonymous sites are not mere silent passengers. Constraints associated with translational efficiency ((Ikemura, 1981, 1985; Plotkin and Kudla, 2011), 1985; , accuracy (Akashi, 1994), mRNA stability (Kudla et al.,2009), and cotranslational protein folding (Kimchi-Sarfaty et al.,2007) impose selective pressures on synonymous substitutions, leading to non-random usage of synonymous codons, a phenomenon known as codon usage bias (CUB). Consequently, the synonymous substitution rate (*dS*) may be systematically depressed in regions under such constraint, leading to an artefactual inflation of *ω* and the potential for erroneous inference of adaptive evolution (Rahman et al., 2021; Verdonk et al., 2025).

Recent investigations into the fitness effects of synonymous mutations have ignited significant debate. For instance, Shen et al. (2022) reported that a substantial majority of synonymous mutations in yeast are strongly non-neutral, with fitness effects comparable to nonsynonymous changes. While these findings have been vigorously contested, with critics arguing that the observed effects may stem from experimental artifacts or specific gene contexts (Kruglyak et al., 2023), the controversy underscores the growing recognition that the “silent” nature of synonymous sites can no longer be assumed *a priori*.

Multiclass Synonymous Substitution (MSS) models address this limitation by partitioning synonymous codons into distinct rate classes, thereby accounting for heterogeneity in the synonymous substitution process. While often motivated by selection for translational efficiency, this framework is sufficiently general to capture systematic rate variation arising from diverse factors, including mutational biases, DNA repair mechanisms such as GC-biased gene conversion (gBGC), and varying DNA repair efficiency across different genomic contexts. In this study, we integrate MSS models into the BUSTED framework for detecting episodic positive selection (Murrell et al., 2015; Kosakovsky Pond et al., 2020). Using simulated and empirical datasets, we demonstrate that unmodelled synonymous selection bedevils standard tests, leading to catastrophic false positive rates in some regimes. The BUSTED+S+MSS approach provides a robust, necessary correction, restoring the validity of evolutionary inference by fundamentally redefining the null hypothesis of synonymous neutrality.

## 2 Methods

### 2.1 Codon evolutionary models

BUSTED (Murrell et al., 2015) tests for evidence that episodic diversifying selection has occurred at a subset of branches and sites in for a given alignment. The statistical test depends on the *ω* parameter, which we have shown to be sensitive to alignment or genome-wide rate differences among synonymous substitutions (Rahman et al., 2021). To mitigate these biases, BUSTED+S incorporates site-specific synonymous rate variation (SRV), which allows the synonymous rate (*α*) to vary across sites according to a discrete distribution. Furthermore, to handle potential alignment artifacts or sequencing errors that can manifest as spurious clusters of substitutions, we utilize an “error-sink” component. This component adds a fourth *ω* class with a very high value (*ω* = 100), designed to absorb sites with biologically implausible rates of nonsynonymous change, thereby preventing them from inflating the test for positive selection (Selberg et al., 2025). Verdonk et al. (2025) developed Multiclass Synonymous Substitution (MSS) models that can account for heterogeneous synonymous substitution rates and correct *ω*. In our current work, we extend BUSTED to incorporate the multiple synonymous substitution rates estimated by MSS models.

The simplest MSS model uses a genetic algorithm to partition synonymous substitutions into two rate classes based on their substitution dynamics. One rate class comprises synonymous codon pairs that exchange at a baseline rate (the “background” or “neutral” class, with shared rate *α*_background_), typically representing exchanges between codons of similar fitness. The other rate class comprises pairs that exchange at a distinct rate (the “selected” or “constrained” class, with shared rate *α*_selected_), capturing exchanges that are inhibited (or promoted) by selective constraints, such as those between optimal and non-optimal codons. BUSTED can also accept more complex MSS models (*e*.*g*., a SynREVCodon model (Verdonk et al., 2025), for which there is one empirically estimated synonymous rate for every exchangeable synonymous codon pair). In both cases, the synonymous rates (or rate classes) provided by the MSS model are incorporated into the BUSTED rate matrix, allowing the instantaneous rate of substitution from 1 sense (*x*) codon to another (*y*) to depend on the identities of the codons being exchanged (Table 1).

**Table 1.** Instantaneous substitution rates from one sense codon (x) to another (y) under BUSTED+MSS.

| One nucleotide? | Synonymous? | Substitution rate | Example |
| --- | --- | --- | --- |
| Yes | Yes | $\alpha_{ij}\theta_{ij}\pi_j^p$ | $\text{AGC} \rightarrow \text{AGT}$:<br>$\alpha_{AGC:AGT}\theta_{CT}\pi_T^3$ |
| Yes | No | $\beta^{bs}\theta_{ij}\pi_j^p$ | $\text{AGC} \rightarrow \text{ACC}$:<br>$\beta^{bs}\theta_{GC}\pi_C^2$ |
| No | * | 0 | $\text{TCC} \rightarrow \text{AGC}$: 0 |

We consider several parameterizations for the synonymous substitution rates *α*(*x, y*), as defined in Verdonk et al. (2025) and summarized in Table 2.

**Table 2.**
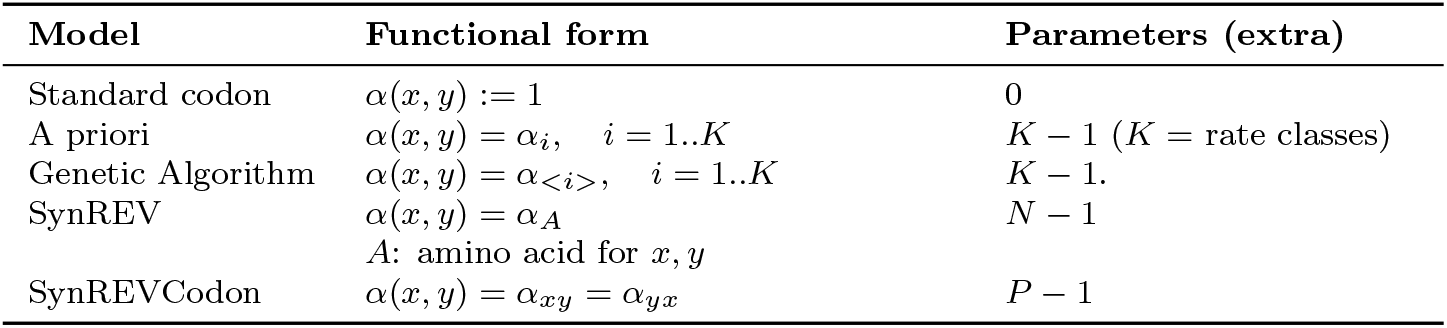
Parameterization of synonymous substitution rates *α*(*x, y*) in different MSS models (Verdonk et al., 2025). *N* denotes the number of amino acids with more than one codon (typically 18), and *P* denotes the number of exchangeable synonymous codon pairs (e.g., *P* = 67 for the universal genetic code).

*θ*_*ij*_ are the nucleotide substitution bias parameters, which follow the convention established by Wisotsky et al. (2020) that *θ*(x, y) = *θ*_nm_ , where **n** and **m** are the 2 nucleotides being exchanged. We assert time-reversibility - *i*.*e*., that *θ*(x, y) = *θ*(y, x) - and assert *θ*_*AG*_ = 1 for identifiability (only products of rates and times can be inferred). 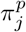 are the position-specific frequencies of each target nucleotide, such that the frequency of a Thymine base in the third codon position is represented by 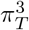. Position-specific nucleotide frequencies are estimated via a corrected empirical F3 × 4 (CF3 × 4) estimator (Pond et al., 2010).

The original BUSTED method (Murrell et al., 2015) models the variation in *ω* using random effects over branches and sites. The BUSTED+S extension (Wisotsky et al., 2020) adds site-to-site (but not branch-to-branch) variation in the synonymous substitution rate *α*_*s*_, drawn from a discrete distribution. BUSTED+S+MSS combines these components with a global codon-pair specific correction *α*_*xy*_ (Table 2; Fig 1). The instantaneous substitution rate from codon *x* to *y* at site *s* is parameterized as proportional to *α*_*s*_ *× α*_*xy*_ for synonymous changes, and *α*_*s*_ *× α*_*xy*_ *× ω* for nonsynonymous changes. For identifiability, we constrain the mean of the global *α*_*xy*_ rates to 1.

**Figure 1.**
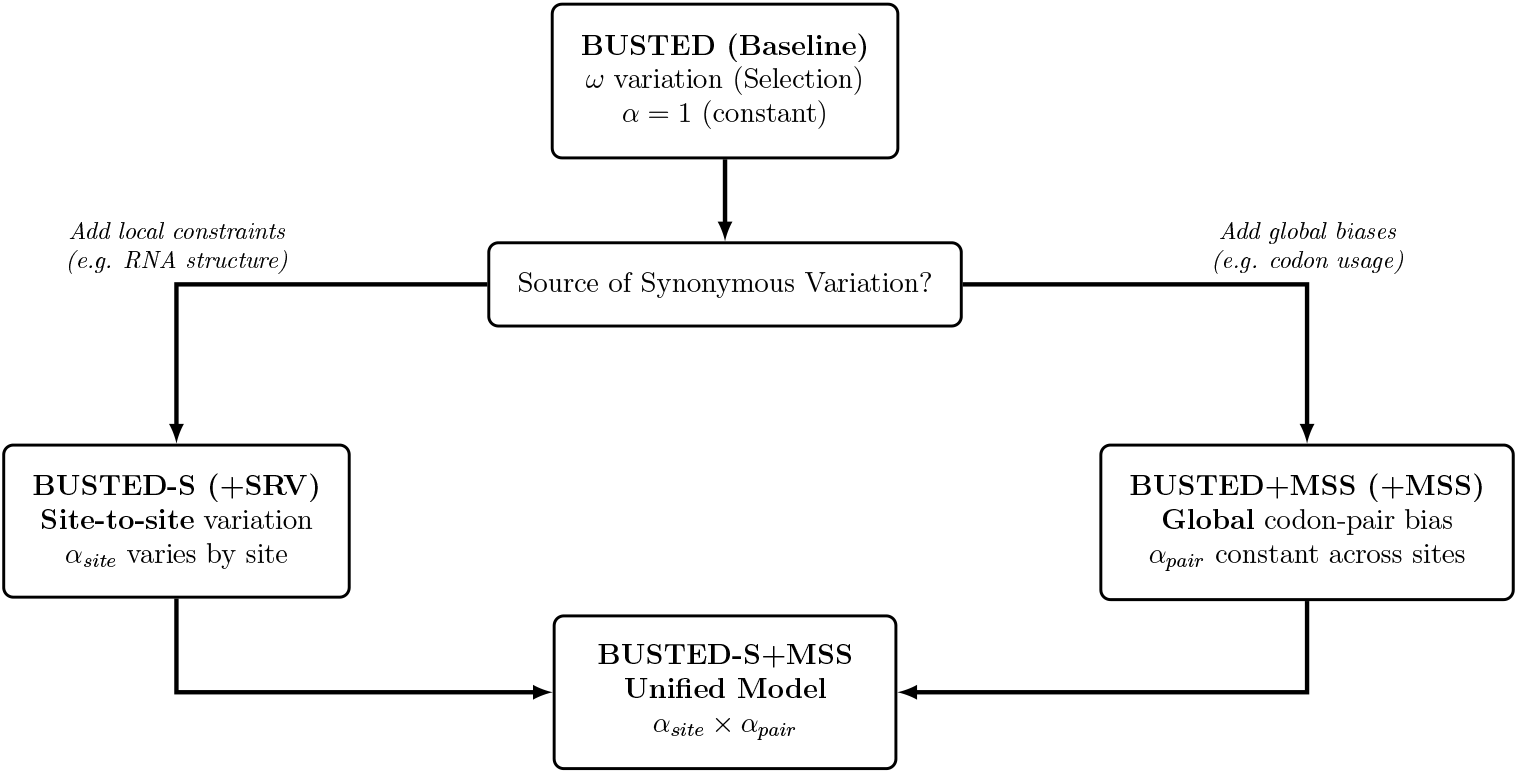
Relationship between BUSTED, BUSTED+S, and BUSTED+MSS models. BUSTED serves as the base-line model where only *ω* varies. BUSTED+S adds site-to-site synonymous rate variation (*α*_*site*_), while BUSTED+MSS adds codon-pair specific rate classes (*α*_*pair*_). The full BUSTED+S+MSS model combines both extensions.

This multiplicative decomposition allows the model to disentangle distinct evolutionary forces. The global codon-pair parameters (*α*_*xy*_) capture genome-wide biases such as those driven by translational efficiency or tRNA abundance, which affect all occurrences of a specific synonymous change. In contrast, the site-specific parameters (*α*_*s*_) capture position-specific evolutionary constraints, such as those imposed by RNA secondary structure or overlapping regulatory motifs, which act locally regardless of the specific codons involved.

The MSS framework accommodates various parameterizations for the synonymous substitution rate *α*(*x, y*), ranging from simple constraints to fully general models (Table 2). The simplest “Standard codon” model assumes a uniform rate (*α*(*x, y*) = 1). *A “priori”* models partition synonymous exchanges into fixed classes derived from external information, such as expression-based codon usage profiles, with a shared rate parameter for each class. “Genetic Algorithm (GA)” models infer these classes directly from the data, optimizing the partition to maximize model fit. More complex parameterizations include “SynREV”, which estimates a distinct rate for each amino acid (assuming all synonymous changes for that amino acid share a rate), and “SynREVCodon”, the most general reversible model, which estimates a unique rate for every pair of exchangeable synonymous codons. In the latter case, the model includes *P −* 1 additional parameters, where *P* represents the number of exchangeable synonymous codon pairs (e.g., *P* = 67 for the universal genetic code).

Finally, we employ the “error sink” extension, BUSTED-E (Selberg et al., 2025), which introduces a specialized rate class with an extremely high synonymous substitution rate. This component is designed to absorb sites with aberrantly high substitution counts that may result from alignment errors or sequencing artifacts, preventing them from inflating estimates of positive selection parameters (*ω*).

### 2.2 Simulation Strategy

To evaluate the potential for synonymous rate variation to confound tests for positive selection, gene alignments without phenotypic selection (*i*.*e*., all *ω* rates were fixed to 1) were simulated under the BUSTED+S+MSS model with the same codon stationary frequencies, mutation rates, and other parameters as a randomly selected *Drosophila* alignment (`FBgn0052113`) and corresponding gene tree. The MSS model used in the simulations was a codon-level partitioned model estimated from `FBgn0052113` via genetic algorithm which partitioned all synonymous one-step substitutions into two classes (background and constrained). Simulating under the BUSTED+S+MSS model allowed the generation of alignments without phenotypic selection with (i) site-to-site synonymous rate variation (*α* multiplier); and (ii) global codon-pair specific rate variation. The MSS model was defined with a background synonymous rate class (*dSn*, set to 1 for identifiability), and a selected synonymous rate class (*dSs*, estimated relative to *dSn*). *dSs* values were chosen along a grid from 0.1 to 1 in steps of 0.05 and 100 alignments without selection (*dN/dS* := 1) were simulated for each *dSs* value, resulting in a total dataset of 1900 alignments. BUSTED+S was then fitted to the simulated alignments with and without the estimated MSS model, and the number of alignments with significant test results under each model was counted. Because all alignments are simulated without selection, each significant result is a false positive, with an expected detection rate of 0.05.

### 2.3 Empirical Dataset Assembly

Representative species were selected from the *Caenorhabditis, Drosophila, Saccharomyces, Enterobacteria*, and primate clades to build the empirical datasets. Each clade has eight or more species with available genome assemblies, and at least one species with strong evidence of selection acting on synonymous substitutions (Ikemura, 1985; Akashi, 1994; Kudla et al., 2009; Trotta, 2013; Simmen, 2008; Kanaya et al., 2001; Marais and Duret, 2001; Plotkin and Kudla, 2011). The dataset of *Drosophila alignments* was prepared as described in Verdonk et al. (2025), and the dataset of Enterobacteria alignments was prepared as described in Rahman et al. (2021).

To build the *Caenorhabditis, Saccharomyces*, and primate datasets, a reference genome for a single representative species per clade was downloaded, along with soft-masked genomes for each other species per clade from the genomes database of the US National Center for Biotechnology Information (NCBI) (Sayers et al., 2022) (Supplemental_S1).

For *Caenorhabditis* and *Saccharomyces*, `cactus` (Armstrong et al., 2020) and the most recent (as of 2022 when the data were collected) unpublished phylogeny of Stevens *et. al* (2024) (*Caenorhabditis*) and the phylogenetic tree of Naseeb *et. al* (Naseeb et al., 2017) (*Saccharomyces*) were used to align all other genomes with respect to the reference genome (specific assembly versions are listed in Supplemental_S1). For primates, `cactus`, the `hal2mafMP.py` script (from the HAL tools suite), and the species tree from Ensembl (Martin et al., 2023) release 109 were used. Gene alignments were then extracted from each whole genome alignment using the coding gene sequence (CDS) coordinates from each reference genome. Any species-specific gene sequences that did not match the reference genome’s strand, exon order, or coding frame were removed, as well as sequences with internal stop codons. Gene alignments with nucleotide lengths that were not a multiple of 3, and alignments with fewer than four aligned sequences, were also excluded.

These clades were chosen to represent a range of population genetic environments and selective pressures on codon usage. *Enterobacteria* and *Saccharomyces* are unicellular organisms with large effective population sizes (*N*_*e*_) and rapid replication cycles, where translational efficiency is a dominant force driving strong codon usage bias (CUB) (Ikemura, 1981, 1985). *Drosophila* and *Caenorhabditis* represent invertebrates with relatively large *N*_*e*_ and well-documented translational selection (Akashi, 1994; Duret, 2002), although other factors like splicing regulation may also play a role (Parmley et al., 2007). In contrast, Primates have much smaller *N*_*e*_, making them less efficient at purging slightly deleterious synonymous mutations; here, synonymous rate variation is expected to be driven less by translational selection and more by mutational biases and GC-biased gene conversion (gBGC) (Duret, 2002; Chamary et al., 2006). In these clades, the MSS model serves as a critical correction not only for selection, but for the systematic mechanistic biases that can otherwise mimic the signal of adaptive evolution.

## 3 Results

### 3.1 Impact of Model Misspecification

We have previously established that selection on synonymous substitutions leads to overestimation of positive selection when the selection on synonymous substitutions is not accounted for (Rahman et al., 2021; Verdonk et al., 2025). However, it is unclear whether accounting for synonymous rate variation across sites (as BUSTED+S does by default) is enough to address differences in substitution rates between different pairs of synonymous codons. Site-to-site variation models (*SRV*) phenomenologically capture scalar differences in evolutionary rate at different positions—reflecting mutation rate variation or site-specific constraints— but they cannot distinguish between different *types* of synonymous changes (e.g., optimal to non-optimal vs. neutral) which occur globally across the gene. To explore this, one random *Drosophila* alignment (`FBgn0052113`) was selected and its parameters (*e*.*g*., codon frequencies, nucleotide mutation rates, etc.) were used to simulate 1900 alignments with no positive selection (*dN/dS* − 1) but with both site-to-site synonymous rate variation and global codon-pair bias. The magnitude of the selected synonymous rate (*dS*_*s*_ = *α*_*sel*_) was systematically varied relative to the neutral synonymous rate (*dS*_*n*_ = *α*_*neutral*_ − 1) in the simulations. BUSTED+S was then fitted to each of the simulated alignments and the number of significant results at *p* ≤ 0.05 was tabulated. Because strictly neutral data was simulated, each detection is a false positive.

A GA-derived two-class MSS model was estimated from the initial *Drosophila* alignment, then BUSTED+S+MSS was re-fitted on the set of 1900 simulated alignments. The false positive rate of BUSTED+S+MSS was compared to BUSTED+S alone (Fig 2). It is observed that even with moderate selection acting on synonymous substitutions (*dS*_*s*_*/dS*_*n*_ = 0.85), failing to account for differences in synonymous substitution rates among synonymous codon pairs leads to a false positive rate of 27%, substantially above the expected 5% threshold. As selection strength increases, this error rate becomes catastrophic (reaching nearly 100% at *dS*_*s*_ = 0.5). BUSTED+S+MSS effectively corrects the false positive rate back to nominal levels (Fig 2).

**Figure 2.**
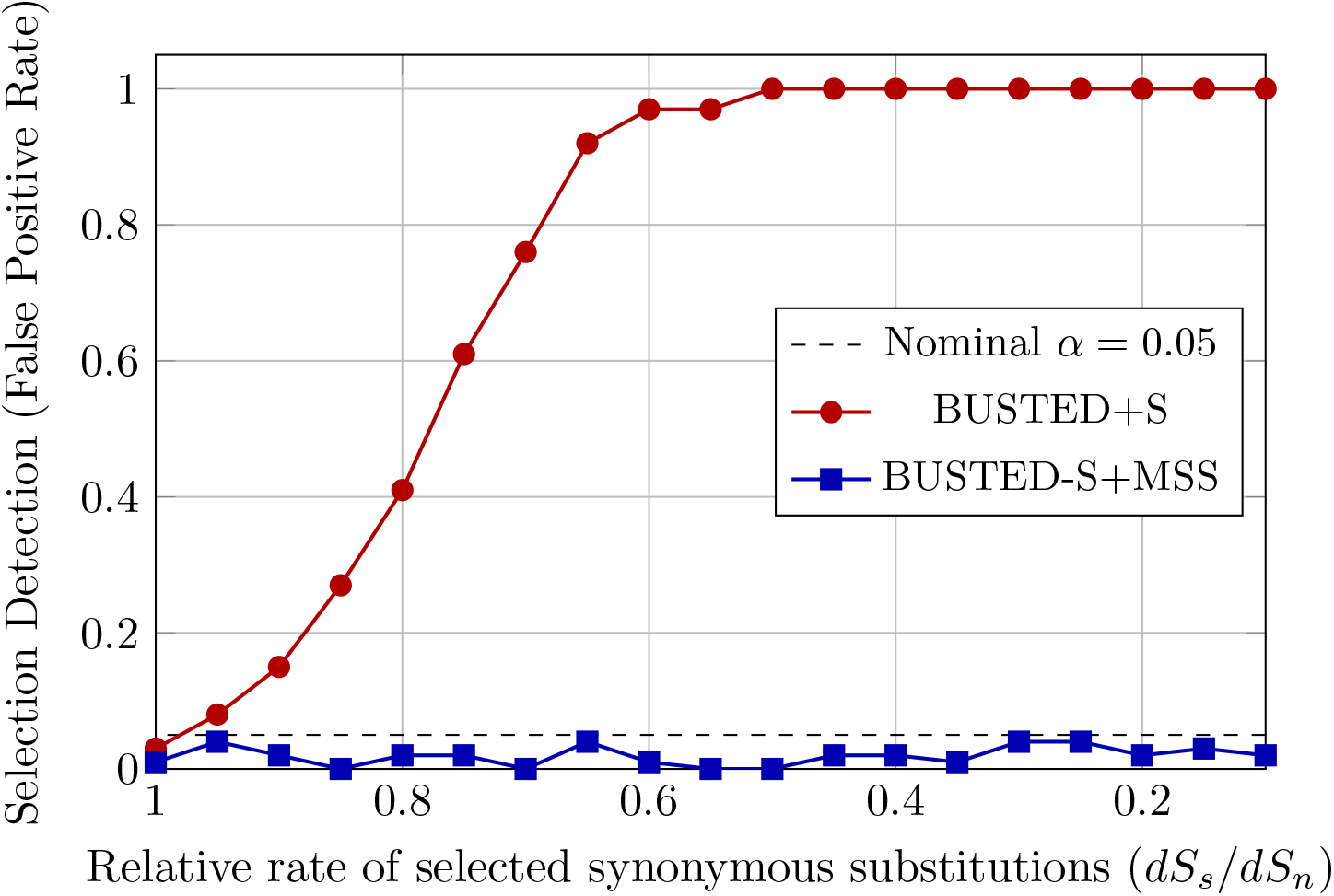
False positive rates (Type I error) under increasing synonymous selection strength. BUSTED+S (red) shows a runaway false positive rate as the synonymous rate of the selected class (*dS*_*s*_) decreases relative to the baseline rate. In contrast, BUSTED+S+MSS (blue) maintains a nominal false positive rate near the expected 0.05 level (dashed black line). *dS*_*s*_ = 1.0 represents unconstrained evolution (no selection).

Furthermore, the BUSTED+S+MSS model provides a significantly better fit to the data as the strength of selection acting on synonymous substitutions increases (Fig S1). When selection is weak or absent (*dS*_*s*_ ≈ 1), the improvement in AIC is negligible or slightly negative (due to parameter penalties), but as selection becomes stronger (lower *dS*_*s*_), the ΔAIC shifts dramatically in favor of the MSS model.

### 3.2 Correction of Selection Inference using MSS

We have previously shown that some of the variation in MSS rates is attributable to selection on cognate tRNA abundance (Verdonk et al., 2025) , which varies across the tree of life (Duret, 2002; dos Reis et al., 2003). However, synonymous substitution rates are also shaped by mutational biases, context-dependent mutation rates, and DNA repair mechanisms (Hershberg and Petrov, 2008). Other well-known selection pressures on synonymous codon usage, like CpG site maintenance (Simmen, 2008), are likely negligible in clades where CpG methylation is nearly absent (Cooper et al., 2005). One would therefore expect that synonymous codon exchangeabilities also differ across clades in response to variation in selective pressure. A single MSS model fit to all clades would “average out” such variation; each clade would require its own MSS model to properly capture that clade’s synonymous substitution rates. This approach is conceptually similar to Dayhoff and colleagues” method for estimating PAM matrices from amino acid substitutions found in closely related proteins (Dayhoff et al., 1978).

MSS models were fitted to 5 diverse empirical datasets (*Caenorhabditis, Drosophila*, Enterobacteria, Primate, and *Saccharomyces*) to investigate how accounting for selection acting on synonymous substitutions affects the detection of positive selection by BUSTED. Each dataset was built from genome-level alignments of at least 8 species, one of which was always a well-studied model species, and contained a suitable number of synonymous substitutions to reliable estimate MSS model parameters (Table 3).

**Table 3.** Overview of empirical datasets. For each dataset, we report the number of gene alignments, median alignment length in codons (Length), median number of taxa (Taxa), median total tree length in substitutions per site (Subs/Site), total count of synonymous substitutions across all genes, and the median number of synonymous substitutions per gene. Ranges are shown in parentheses.

| Dataset | Alignments | Length | Taxa | Subs/Site | Total Syn. Subs | Syn. Subs/Gene |
| --- | --- | --- | --- | --- | --- | --- |
| *Caenorhabditis* | 7,696 | 335 (100–4,858) | 8 (4–46) | 1.2 (0.11–4.2) | 3,749,028 | 386 (3–6,281) |
| *Drosophila* | 12,062 | 411 (21–8,933) | 12 (4–23) | 0.48 (0.017–2.0) | 4,215,193 | 255 (0–5,762) |
| *Enterobacteria* | 1,594 | 332 (66–1,523) | 13 (4–15) | 2.0 (0.17–8.4e3) | 1,457,215 | 812 (0–4,022) |
| Primate | 14,371 | 466 (44–5,137) | 25 (8–26) | 0.40 (0.045–1.9) | 5,735,422 | 292 (3–5,557) |
| *Saccharomyces* | 4,916 | 418 (100–4,092) | 7 (4–16) | 0.63 (0.023–2.1) | 1,533,103 | 263 (8–2,077) |

For each clade, a genetic algorithm (GA) was employed to “infer” the specific pattern of synonymous rate variation from the data, without *a priori* assumptions about the driving mechanism (*e*.*g*., tRNA vs. methylation or mutational bias). Consistent with our simulation study, an optimal partition of synonymous codon pairs into two rate classes: a “background” class (baseline rate) and a “constrained” class (reduced rate) was sought. This training was performed on a random subsample of 500 gene alignments per dataset to avoid overfitting and ensure computational feasibility. The resulting consensus partition of codon pairs (into “background” and “constrained”), which minimizes the BIC across the training set, was then fixed and used to define a clade-specific MSS model. This model included one more parameter compared to the baseline BUSTED+S, the *dS*_*s*_*/dS*_*n*_ ratio, which was estimated from each gene as part of the process of testing for positive selection.

While theoretically appealing, estimating a complex MSS model like SynREVCodon for every individual gene is often statistically underpowered, particularly for short alignments with sparse synonymous substitutions. This risks overfitting, where the model captures stochastic noise rather than biological signal. Conversely, assuming a single global model for an entire clade ignores the gene-specific variation in selection intensity. Our approach strikes a balance: we infer the *structure* of synonymous rate variation (the partitioning of codons into classes) at the clade level, leveraging the power of hundreds of alignments. This yields a robust, empirically grounded model with few parameters to be estimated for each individual gene. This strategy is both statistically rigorous—avoiding parameter bloat—and computationally efficient, enabling genome-wide scans that would be prohibitive with per-gene full model inference.

The derived MSS models exhibit substantial heterogeneity in selection patterns across the five clades (Fig 3). Whilst some synonymous codon exchanges are universally identified as either background or constrained, a significant proportion exhibit clade-specific rate classifications. This variability likely reflects the differing contributions of translational efficiency, mutational biases, and other constraints unique to each taxonomic group. The presence of these idiosyncratic patterns reinforces the necessity of the empirical, data-driven approach employed in this study to ensure that clade-specific synonymous constraints are accurately accounted for during selection inference.

**Figure 3.**
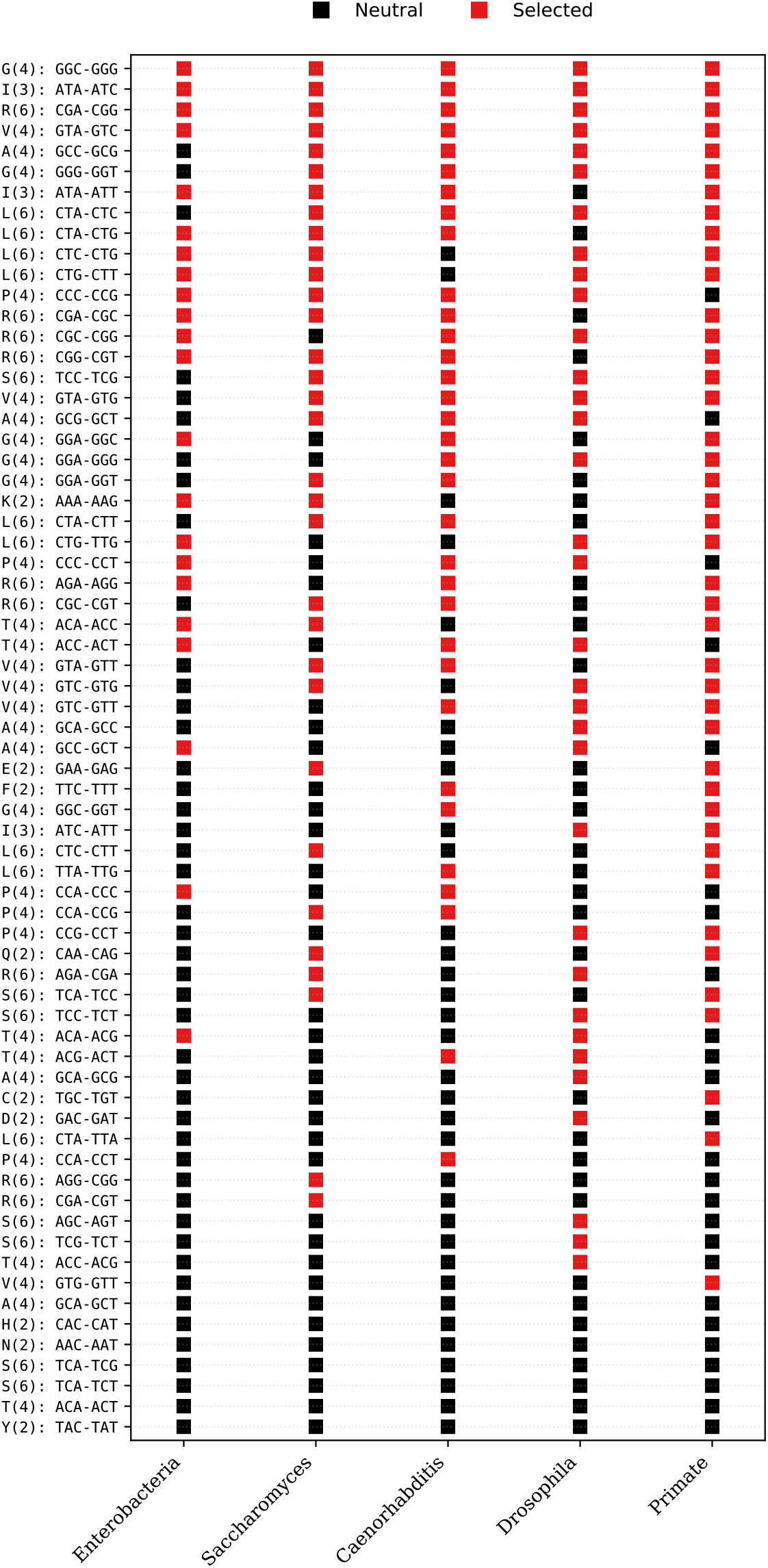
Visualization of the clade-specific MSS models derived from empirical data. Each row represents a specific synonymous codon exchange grouped by amino acid and fold redundancy (e.g., S(6): TCG-TCT). Each column represents one of the five taxonomic clades analyzed. Squares are colored by the inferred rate class: Black for Background (baseline synonymous rate), Red for Selected (distinct synonymous rate). Pairs are sorted by selection frequency across clades (ascending) and secondarily by amino acid (reverse-alphabetically, descending). This sorting places universally selected pairs and those belonging to alphabetically earlier amino acids at the top of the visualization. The variation in patterns across columns demonstrates that selection pressures on codon usage are clade-specific, necessitating empirical estimation for accurate correction.

BUSTED+S was fitted (`--starting-points 5 --srv Yes --error-sink Yes`) to each dataset with and without the corresponding MSS model in order to test for evidence of selection and compare model fit for each gene. Fewer genes were found with significant evidence of positive selection across all datasets when using the BUSTED+S+MSS model, compared to the BUSTED+S model alone. The BUSTED+S+MSS model is also a better fit across all datasets, with as much as a 55 point improvement in mean AIC score (Table 4). Notably, in the Primate dataset, the number of genes identified as evolving under positive selection was reduced by over 50% (from 196 to 84 unique detections), highlighting that mechanistic biases such as gBGC can drive false positives as effectively as translational selection. Similarly, in *Saccharomyces*, the low number of genes initially identified as significant (26, or 0.5%) is consistent with the strong global purifying selection characteristic of this genus; the further reduction to just 4 unique detections under MSS underscores that even these rare signals can be confounded by synonymous constraints.

**Table 4.** Fitting BUSTED to empirical datasets. We report the number of genes (N) with evidence of positive selection (*p* ≤ 0.05), selection intensity (*ω*_3_), and fraction of selected sites (*p*_3_) for three mutually exclusive categories: genes significant only in BUSTED+S (parameters from BUSTED+S), genes significant only in BUSTED+S+MSS (parameters from BUSTED+S+MSS), and genes significant in both models (parameters from BUSTED+S+MSS). Values in parentheses in the “N” column represent the percentage of total genes analyzed. ΔAICc shows the average improvement in fit (BUSTED+S AIC - BUSTED+S+MSS AIC). Outliers (*ω*_3_ *>* 100) are excluded from means.

| Dataset | BUSTED+S Only |  |  |  | BUSTED+S+MSS Only |  |  |  | Both |  |  |  |
| --- | --- | --- | --- | --- | --- | --- | --- | --- | --- | --- | --- | --- |
| | N | $\omega_3$ | $p_3$ | $\Delta AICc$ | N | $\omega_3$ | $p_3$ | $\Delta AICc$ | N | $\omega_3$ | $p_3$ | $\Delta AICc$ |
| *Caenorhabditis* | 155<br>(2.0%) | 6.55<br>(2-77) | 0.017<br>(0-0.13) | 52.1<br>(-22-380) | 34<br>(0.4%) | 8.16<br>(4-24) | 0.011<br>(0-0.05) | 42.5<br>(-1-325) | 143<br>(1.9%) | 6.19<br>(2-39) | 0.020<br>(0-0.56) | 33.6<br>(-2-572) |
| *Drosophila* | 144<br>(1.2%) | 5.33<br>(2-25) | 0.064<br>(0-0.71) | 13.1<br>(-5-264) | 70<br>(0.6%) | 7.56<br>(2-31) | 0.024<br>(0-0.24) | 14.0<br>(-2-149) | 315<br>(2.6%) | 7.45<br>(2-63) | 0.041<br>(0-1) | 7.4<br>(-3-261) |
| *Enterobacteria* | 23<br>(1.4%) | 8.98<br>(3-35) | 0.011<br>(0-0.03) | 60.5<br>(0-128) | 12<br>(0.8%) | 8.20<br>(4-23) | 0.009<br>(0-0.01) | 53.3<br>(8-131) | 17<br>(1.1%) | 11.10<br>(3-99) | 0.012<br>(0-0.03) | 63.3<br>(-2-184) |
| Primate | 196<br>(1.4%) | 8.72<br>(1-72) | 0.066<br>(0-0.53) | 23.2<br>(-9-165) | 84<br>(0.6%) | 18.94<br>(2-83) | 0.011<br>(0-0.38) | 22.0<br>(-6-139) | 322<br>(2.2%) | 9.51<br>(1-92) | 0.039<br>(0-0.46) | 19.6<br>(-8-260) |
| *Saccharomyces* | 26<br>(0.5%) | 5.99<br>(2-35) | 0.021<br>(0-0.12) | 8.4<br>(-1-28) | 4<br>(0.1%) | 8.95<br>(3-16) | 0.016<br>(0-0.02) | 8.2<br>(-1-22) | 36<br>(0.7%) | 6.61<br>(3-12) | 0.057<br>(0-1) | 4.0<br>(-2-19) |

The empirical estimates of *dSs* in *Enterobacteria* underscore the prevalence of non-neutral synonymous evolution (Fig 4). We observed a clear divergence in selection intensity associated with model preference: genes for which BUSTED+S+MSS provided a superior fit exhibited significantly stronger purifying selection acting on synonymous substitutions (mean *dSs* ≈ 0.41) than those where the MSS model was not preferred (mean *dSs* ≈ 0.82; Mann-Whitney U test, *p<* 10^*−*22^). This strong association indicates that the statistical preference for the MSS framework is directly driven by the magnitude of the synonymous constraint.

**Figure 4.**
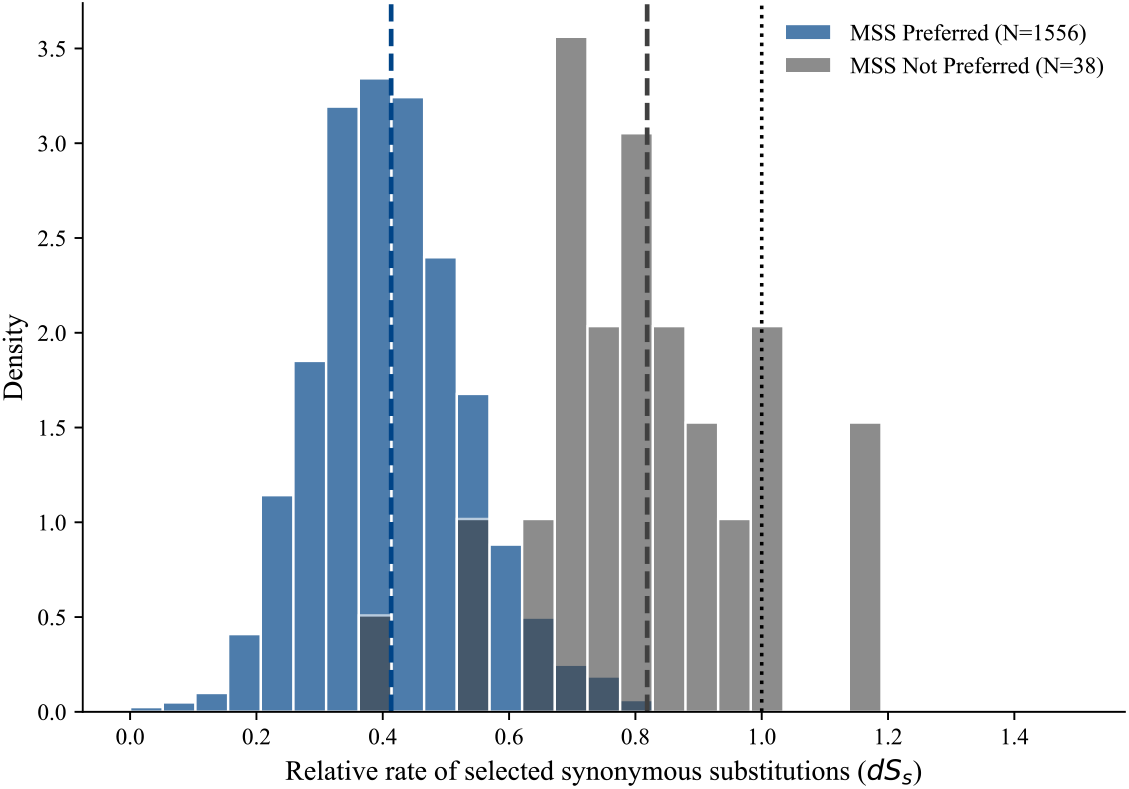
Distribution of the relative rate of selected synonymous substitutions (*dSs*) in *Enterobacteria*, stratified by model preference. Genes where the MSS model provides a better fit (ΔAIC > 0, blue) exhibit significantly stronger purifying selection acting on synonymous substitutions (mean *dSs* ≈ 0.41) compared to genes where the MSS model is not preferred (gray, mean *dSs* ≈ 0.82; Mann-Whitney U test, *p <* 10^*−*22^). This confirms that the statistical preference for the MSS model is directly driven by the presence of non-neutral synonymous evolution.

MSS model inclusion also impacted key parameter estimates such as *ω*_3_, which reflects the selection intensity of all branch/site pairs under positive selection, and *p*_3_, the fraction of sites under positive selection. The mean *ω*_3_ estimated under BUSTED+S+MSS is larger by a factor of 1.5 in the *Drosophila* dataset and by a factor of 1.4 in the *Enterobacteria* dataset when compared to the mean *ω*_3_ estimated under BUSTED+S. In contrast, the mean *ω*_3_ estimated under BUSTED+S+MSS is slightly reduced compared to the mean BUSTED+S estimate in the *Caenorhabditis, Saccharomyces*, and primate datasets. *p*_3_ was reduced across all datasets under BUSTED+S+MSS, by as much as a factor of 1.8 in the *Drosophila* dataset.

The practical impact of model choice is illustrated by *CG11912* (FBgn0031248), a *Drosophila* gene encoding a predicted serine hydrolase involved in proteolysis. This gene exhibits moderately biased codon usage (*CAI* = 0.400, calculated using the cai tool and `Edrome_high.cut` reference set in (Rice et al., 2000)). When analyzed with the standard BUSTED+S model, *CG11912* shows statistically significant evidence of positive selection (*p* = 0.042), with a subset of sites (*p*_3_ ≈ 21%) evolving at an elevated rate (*ω*_3_ ≈ 2.2). However, this signal disappears when accounting for global selection acting on synonymous substitutions: the BUSTED+S+MSS model finds no evidence of selection (*p* = 0.439), estimating a much lower *ω*_3_ ≈ 1.2. The inferred selected synonymous rate for this gene was *dS*_*s*_ ≈ 0.7, indicating purifying selection acting on synonymous substitutions.

#### 3.2.1 Analysis of Model Discordance

To better understand the biological impact of accounting for MSS, the subset of genes where BUSTED+S and BUSTED+S+MSS yielded discordant results was examined. The most common case of discordance occurred when BUSTED+S initially detected positive selection, but the signal was lost upon including the MSS model. These genes almost universally preferred the BUSTED+S+MSS model, with mean AICc improvements comparable to those seen across the full datasets. In these genes, the inferred selection intensity (*ω*_3_) was dramatically reduced under BUSTED+S+MSS (by 23–54% compared to BUSTED+S), while the fraction of sites under selection (*p*_3_) remained largely unchanged. This pattern suggests that unmodeled selection acting on synonymous substitutions often manifests as an inflated *ω*_3_, which BUSTED+S+MSS correctly reassigns to the synonymous substitution component.

Conversely, a smaller number of genes only reached statistical significance after accounting for MSS. These genes also preferred the BUSTED+S+MSS model. In these cases, the effect on *ω*_3_ was more variable across clades, but often resulted in an increase in estimated selection intensity (Supplementary Table S2). This indicates that in some genes, synonymous rate variation can mask true signals of positive selection by “diluting” the relative impact of nonsynonymous changes.

A robust core of genes retained significant evidence of positive selection under both models (Table 4, “Both” column). For these genes, the BUSTED+S+MSS model estimates of selection intensity (*ω*_3_) were generally high, reinforcing that the MSS correction does not simply abolish all signals of selection, but rather refines them by removing false positives driven by codon usage bias. Consequently, positive selection detected by BUSTED+S+MSS can be interpreted with greater confidence as reflecting adaptive amino acid evolution, decoupled from the confounding effects of synonymous constraint.

The drivers of discordance in the *Enterobacteria* dataset were further investigated by analyzing the estimated strength of selection acting on synonymous substitutions (*dSs*). It was found that genes which lost their positive selection signal under the MSS model had significantly lower *dSs* values (mean 0.40) compared to those that retained significance (mean 0.52). This relationship remained significant (*p* = 0.024) even after controlling for the strength of the original evidence (LRT) and model fit improvement in a multivariate logistic regression. This confirms that strong purifying selection acting on synonymous substitutions (*dSs«* 1), if unmodeled, artificially depresses the baseline substitution rate, leading standard models to misinterpret the relative nonsynonymous rate as adaptive evolution.

To identify the factors that best predict whether a gene initially classified as significant under BUSTED+S will lose its significance under BUSTED+S+MSS, a logistic regression analysis was performed for each dataset. As expected, the strength of the original evidence was the primary determinant: genes with higher likelihood ratio statistics were less likely to lose their positive selection status (*p <* 0.01 for all datasets). However, even after controlling for initial signal strength, the improvement in model fit provided by the MSS model (ΔAICc) was a significant predictor of signal loss in *Drosophila* (*p* = 0.037) and was the strongest predictor in *Caenorhabditis* (*p <* 10^*−*5^), and marginally significant for *Saccharomyces* (*p* = 0.068). This confirms that for genes where global codon-pair bias is a prominent feature, standard codon models are systematically prone to false positives. In contrast, this effect was not statistically significant in Primates or *Enterobacteria*, which may reflect different underlying evolutionary dynamics or the smaller number of significant genes available for analysis in these clades.

#### 3.2.2 Interplay between Synonymous Rate Variation and Codon Bias

Factors including selection to maintain gene function (Parmley et al., 2006) , and mRNA secondary structure (Kudla et al., 2009) alter the synonymous substitution rate across alignment sites somewhat independently of codon preference, a phenomenon known as synonymous rate variation (SRV). BUSTED accounts for the potential presence of SRV by default when testing for selection (Wisotsky et al., 2020). Given that both SRV and MSS model the effects of selection on synonymous substitution rates (positional vs. codon-specific), we investigated the extent to which these two components capture distinct versus overlapping evolutionary signals.

The *Enterobacteria* dataset, which showed the strongest preference for MSS models (Table 4), was analyzed. The relationship between the two model components was formally assessed by defining an interaction term based on model fit improvement. We asked: does adding MSS to a model that already has SRV provide any additional benefit?

Analysis of the *Enterobacteria* dataset revealed that while the two models share some information (redundancy), they are largely complementary. For the vast majority of genes (81.4%), the combined BUSTED+S+MSS model provided a significantly better fit than either component in isolation. Specifically, the combined model yielded a mean log-likelihood improvement of 161.22 units over the baseline, substantially higher than the improvement from either SRV (132.60) or MSS (35.42) alone. This confirms that site-specific constraints (SRV) and global selection acting on synonymous substitutions (MSS) capture distinct biological signals, and both are essential for accurately modeling synonymous evolution.

To further visualize the relative contributions of each component, we partitioned the total model improvement for each gene into unique, redundant, and synergistic shares (Fig 5). For the majority of genes (redundant case), the distribution reveals that site-to-site rate variation (Unique SRV) is the dominant driver of model fit improvement, though unique codon-pair bias remains a consistent factor. In contrast, the subset of synergistic genes displays a distinct pattern where the combined model unlocks additional explanatory power beyond the sum of the individual components. This synergy challenges the conventional view that synonymous constraints are merely additive parameters. Instead, it reveals a biological interaction: correctly modeling global codon preferences de-noises the baseline rate estimates, thereby unmasking site-specific rate variation that was previously obscured. This implies that ‘neutral’ synonymous sites are not just rare, but context-dependent.

**Figure 5.**
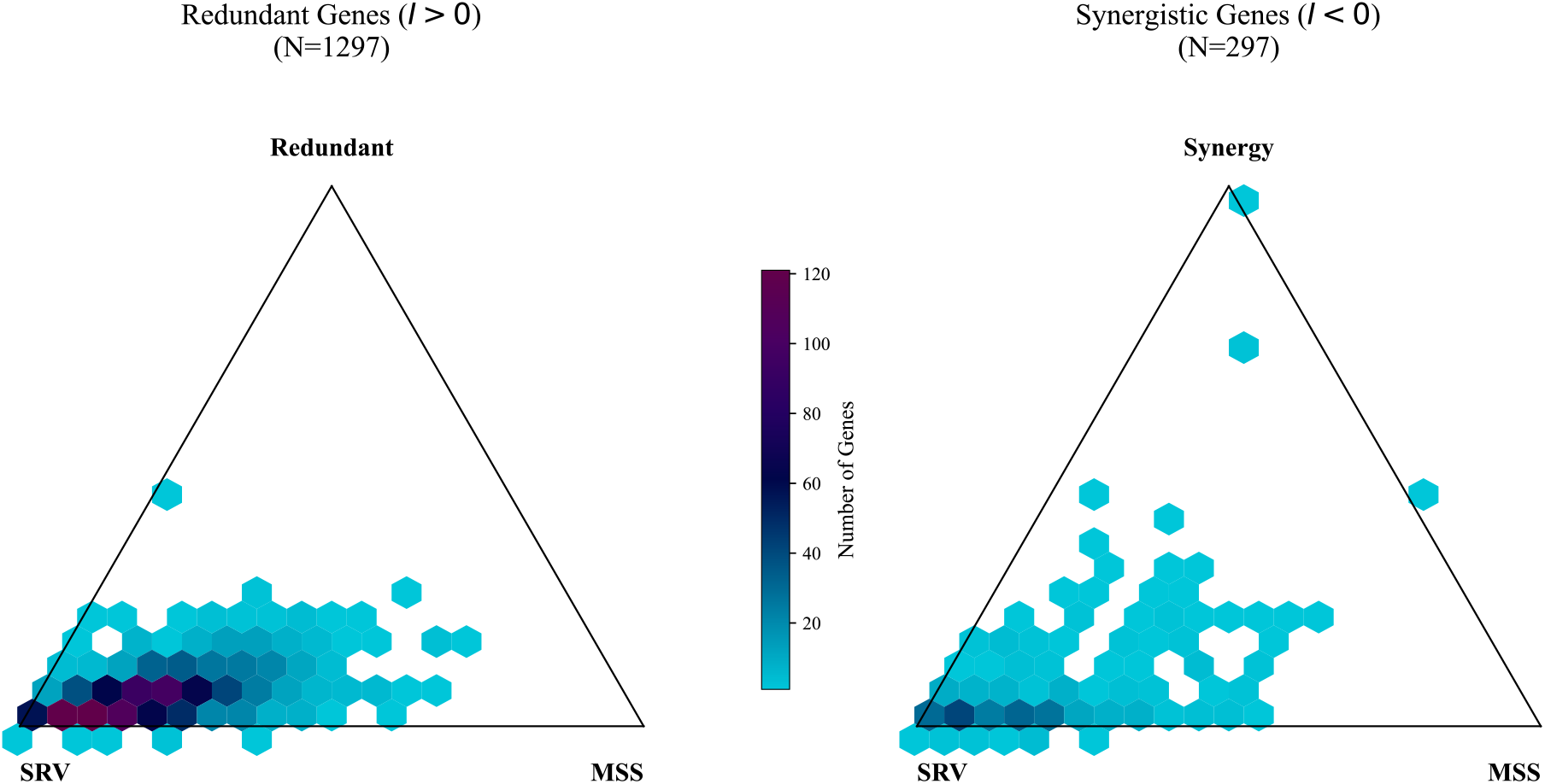
Ternary decomposition of model fit improvement for *Enterobacteria* genes. **(Left)** Genes with redundant signals (*I >* 0). The total improvement is partitioned into the SRV component (Unique SRV), the MSS component (Unique MSS), and the Redundant component. **(Right)** Genes with synergistic signals (*I <* 0). The total improvement is partitioned into the SRV component, the MSS component, and the Synergy term. The heatmap density shows that for the redundant majority, improvement is driven primarily by the SRV component, whereas synergistic genes often show balanced contributions from both factors plus a synergy term.

Beyond the general correlation between expression level and conservation, specific molecular mechanisms likely drive this overlap. For instance, local mRNA secondary structures, which can be modeled as site-specific constraints (SRV), often influence translation initiation and elongation rates (Kudla et al., 2009). Since codon usage bias also regulates translation speed (Tuller et al., 2010), selection may act on both simultaneously to optimize local ribosome dynamics, creating a signal that is detectable by both model components. To determine the independent predictors of this synergy, a logistic regression analysis was performed. Alignment length (*β* = −0.0023,*p <* 0.001) and the number of taxa (*β* = −0.14,*p <* 0.001) were found to be significant negative predictors, confirming that data sparsity is a primary driver of the observed synergy. Conversely, total tree length (*β* = 0.24,*p <* 0.001) was a positive predictor, suggesting that higher sequence divergence also contributes to the phenomenon, possibly by introducing complex substitution patterns that require joint modeling. GC content was also identified as a significant negative predictor (*β* = −7.85,*p* = 0.001), while the intensity of selection acting on synonymous substitutions (*dSs*) was not significant after controlling for other factors. This profile is consistent with the hypothesis that these genes may include horizontally transferred elements (HGT, which often deviate from genomic codon consensus) or rapidly evolving sequences where standard model assumptions are most challenged.

### 3.3 Best Practices for Selection Analysis

The following recommendations are proposed to ensure the robustness of selection analyses in the presence of complex synonymous substitution patterns.

1. **Baseline Analysis:** Begin with the BUSTED+S model (`–srv Yes`). This accounts for site-to-site synonymous rate variation, which is the most pervasive confounder in selection tests.
2. **Evaluating Codon Bias:** If the dataset involves highly expressed genes, compact genomes (e.g., bacteria, viruses), or exhibits strong purifying selection (*dN/dS«* 1), the presence of global codon usage bias should be assessed. For the clades analyzed in this study (e.g., *Drosophila, Enterobacteria*), pre-computed MSS models are provided in the Supplementary Material and can be directly applied. For other lineages, select 20–50 representative alignments—ensuring they are free of gross misalignment— and derive a clade-specific MSS model using the genetic algorithm. Detailed technical instructions for this derivation are provided in the Supplementary Material.
3. **Model Selection:** Compare the fit (AICc) of BUSTED+S versus BUSTED+S+MSS on this subset. If the MSS model consistently provides a superior fit (ΔAICc *>* 10), it should be applied to the full dataset to correct for synonymous constraints. Note that while training the model requires multiple alignments, the test itself is applicable to small datasets (min. 5–8 taxa), provided sufficient divergence exists.
4. **Quality Control:** Always employ the error-sink model (`–error-sink Yes`) to mitigate false positives driven by sequencing errors or alignment artifacts (Selberg et al., 2025). Additionally, screen for recombination (*e*.*g*., using GARD); BUSTED can accommodate the resulting non-recombinant partitions, ensuring that topological incongruence does not confound the test for selection.

## 4 Discussion

In this manuscript, we have integrated a codon model capable of accounting for selection acting on synonymous substitutions (MSS) into an existing statistical test for positive selection (BUSTED (Murrell et al., 2015)). Such selection, if unaddressed, introduces significant false positive rates in tests for adaptive evolution. We demonstrate that MSS models mitigate these errors by explicitly modelling constraints on codon usage. Furthermore, BUSTED+S+MSS consistently provides a superior fit to empirical alignments compared to BUSTED+S, indicating that the MSS component captures biological signal distinct from site-specific rate variation.

The ubiquity of synonymous selection observed across these diverse lineages underscores a fundamental biological reality: the coding sequence is a multi-layered information storage device. The strong correlation between model preference and synonymous constraint in *Enterobacteria* suggests that for a significant fraction of the genome, the maintenance of translational kinetics or mRNA structural integrity imposes a significant selective burden, even if it is generally less intense than the constraints on protein function. In other clades such as *Caenorhabditis*, these pressures are likely compounded by selection for splicing efficiency and the maintenance of Exonic Splicing Enhancers (ESEs). While this signal is likely driven by highly expressed genes, the BUSTED+S+MSS framework is agnostic to the underlying mechanism. Ideally, models could be trained on specific gene subsets to capture expression-dependent constraints, but the primary benefit of the method lies in accounting for the existence of rate heterogeneity itself, which significantly improves the robustness of selection inference. Consequently, the assumption of synonymous neutrality—long the bedrock of molecular evolution—must be discarded in favour of models that explicitly parameterize this “silent” constraint.

It must be noted that the BUSTED+S+MSS framework relies on the availability of a representative set of alignments to infer the baseline synonymous rate classes. The resulting clade-specific model effectively represents a genome-wide average of synonymous constraints; however, in many organisms, the intensity and even the direction of synonymous selection can vary with gene expression levels (Duret, 2002). In lineages with sparse genomic data or extreme divergence, the derivation of a robust MSS model may be challenging. Furthermore, while the current implementation partitions codons into discrete rate classes, the underlying biological reality is likely a continuous fitness landscape. Future extensions could incorporate continuous covariates, such as gene expression levels or tRNA abundance, directly into the rate matrix to provide a more granular correction.

Our approach occupies a distinct niche in the landscape of evolutionary models. Unlike previous attempts that treated synonymous rate variation as a nuisance parameter or required prohibitive computational resources, BUSTED+S+MSS provides the first scalable, generalizable framework to correct for global synonymous constraints within an episodic selection test. This is not merely a refinement of parameter estimates, but a fundamental restructuring of the null hypothesis: from ‘synonymous sites are neutral’ to ‘synonymous sites reflect a baseline of background and constrained evolution.’ Mechanistic frameworks, such as the Halpern-Bruno (Halpern and Bruno, 1998) and swMutSel (Tamuri et al., 2012) models, explicitly parameterize the fitness of each codon to characterize site-specific constraints. While BUSTED+S+MSS shares fundamental assumptions with these models, notably time-reversibility and stationarity of the substitution process, it is optimized for a different inferential objective. MutSel models are effectively the preferred tool for mapping static fitness landscapes and characterizing long-term purifying selection. In contrast, BUSTED+S+MSS is designed to identify episodic bursts of adaptive evolution across branches and sites by allowing the selection parameter (*ω*) to vary. By incorporating a data-driven correction for global synonymous rate variation, our framework preserves this episodic sensitivity whilst mitigating the baseline rate biases that bedevil simpler models, offering a robust solution for hypothesis testing in the presence of complex synonymous constraint.

While BUSTED+S+MSS offers a robust correction, it is not without limitations. Although the inclusion of additional parameters to model synonymous constraint can theoretically reduce power to detect weak selection in small datasets (increasing Type II error), evaluations of analogous extensions to the BUSTED framework—such as those accounting for site-to-site synonymous rate variation (Wisotsky et al., 2020) or multiple nucleotide mutations (Lucaci et al., 2021)—have shown that such corrections generally result in a favorable trade-off between power and false positive rate control. However, the accuracy of the clade-specific MSS model depends on the availability of a representative training set (ideally 20–50 alignments); in lineages with sparse genomic data, the inferred rate classes may be unstable. Additionally, the method assumes that the partition of synonymous codons into “background” and “constrained” classes is conserved across the clade. This assumption may be violated by lineage-specific shifts in effective population size (*e*.*g*., transitions between selfing and outcrossing mating systems), major changes in tRNA pools, or variations in GC-biased gene conversion intensity, potentially necessitating gene-specific model derivation. Similarly, the current MSS implementation does not explicitly model neighborhood-dependent substitution processes, such as the hypermutability of CpG dinucleotides in mammals. While these effects are partially absorbed into the aggregate rate classes, they represent a distinct layer of complexity not fully captured by codon-independent models. Furthermore, unlike the granular treatment of synonymous sites, our current framework does not partition nonsynonymous substitutions into distinct classes (*e.g*., conservative vs. radical). Consequently, heterogeneity in amino acid exchangeability is absorbed into the aggregate *ω* parameter, potentially obscuring finer-scale selective signals. Ultimately, the integration of synonymous selection into standard testing frameworks represents a critical step towards more biologically realistic models of sequence evolution, paving the way for more accurate and nuanced investigations into the molecular basis of adaptation.

## Supporting information

All Supplemental Material

## 5 Data and Code Availability

All empirical datasets, pre-computed MSS models, and custom analysis scripts used in this study are available https://github.com/hverdonk/BUSTED_MSS_Supplemental. able at is implemented as part of the HyPhy software package (Pond et al., 2005; Kosakovsky Pond et al., 2020),The BUSTED+S+MSS framework , Kosakovsky Pond et al., 2020), and can be accessed via the `busted` standard analysis. The Genetic Algorithm for deriving custom MSS models is available through the `MSS-GA` analysis in HyPhy.

## Notes

### Competing Interest Statement

The authors have declared no competing interest.

### Funding

- US National Institute of General Medical Sciences [NIGMS]: GM151683, GM144468
- US National Institute of Allergy and Infectious Diseases [NIAID]: AI183870

