## Supplementary material for "Correcting for Global Synonymous Selection Improves the Accuracy of Episodic Positive Selection Inference": All Supplemental Material

### 6 Supplementary Material

#### S1: Derivation of Clade-Specific Models

The MSS models generated for this study are available in the Supplementary Material (https://github.com/hverdonk/BUSTED_MSS_Supplemental). Researchers can also derive custom clade-specific MSS models for their own datasets. We recommend using at least 20 representative alignments to ensure robust parameter estimation. Ideally, these alignments should be drawn from the specific taxonomic group being analyzed (*e.g.*, Genus or Family) to ensure that the inferred synonymous constraints accurately reflect the lineage-specific biases. A high-quality training set should include genes with sufficient synonymous divergence to inform rate estimation, but avoid saturation. Crucially, the training set must be representative of the clade's "synonymous phenotype," as the method assumes that the partition of synonymous substitutions into "background" and "constrained" classes is conserved across the training genes. Including an excessive number of genes (*e.g.*, >1000) is generally not productive, as the model parameters typically converge with fewer alignments while computational runtimes degrade significantly (*e.g.*, scaling linearly with the number of alignments).

The Genetic Algorithm (GA) employed here explores the space of possible synonymous rate partitions. It optimizes the assignment of the one-step (exchangeable) synonymous codon pairs (*e.g.*, 67 for the universal genetic code) into discrete rate classes (typically $K = 2$) by maximizing the joint c-AIC (or Bayesian Information Criterion, BIC) across the training set; complete implementation details are provided in Verdonk et al. (2025). This stochastic search effectively handles the combinatorial complexity of the model space, identifying robust patterns of shared synonymous constraint without examining every possible permutation.

To infer an MSS model, provide a list of alignment file paths to the genetic algorithm:

```
mpirun -np {threads} HYPHYMPI MSS-GA \
    --ic BIC \
    --mss-type SynREVCodon \
    --classes 2 \
    --filelist your_alignments.txt \
    --output outfile.json
```

The resulting `outfile.json` contains the optimal codon partition. This can be converted into a BUSTED-compatible format using the processor tool:

```
hyphy MSS-GA-processor \
    --json outfile.json \
    --tsv outfile.tsv
```

Finally, the custom MSS model can be applied to BUSTED+S analyses:

```
hyphy busted \
    --alignment gene.nex \
    --mss Yes \
    --mss-type Codon-file \
    --mss-file outfile.tsv \
    --mss-neutral NEUTRAL
```

For most applications, the two-class GA model provides an optimal balance between biological realism and computational efficiency. We recommend verifying the quality of alignments using automated tools like `hyphy-analyses/codon-msa` prior to model derivation, as alignment errors can confound the genetic algorithm's ability to partition synonymous substitution rates accurately.

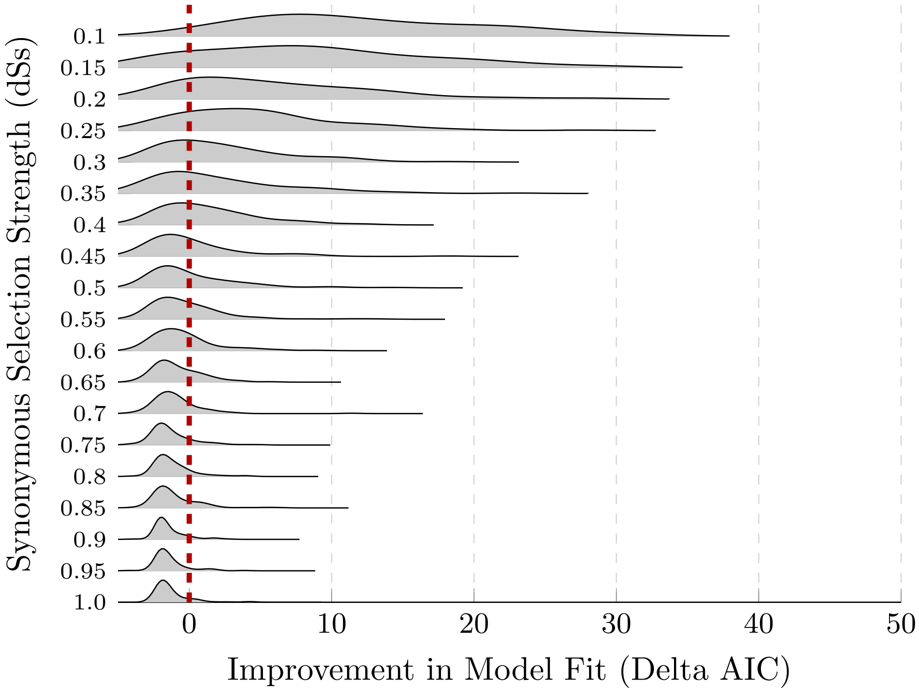


**Figure S1:** Distributions of model fit improvement ($\Delta$AIC = AIC$_{BUSTED+S}$ - AIC$_{BUSTED+S+MSS}$) across 100 replicates for varying synonymous selection strengths ($dS_s$). As selection strength increases (lower $dS_s$, moving up the y-axis), the MSS model provides an increasingly superior fit to the data compared to the baseline BUSTED+S model.
